# Exploring Genome-Wide Mutation Dynamics and Bacterial Cellulose Impairment in *Komagataeibacter intermedius* Cultivated Under Agitation Stress

**DOI:** 10.1101/2025.10.09.681338

**Authors:** Davide Bersanetti, Rahul Mangayil

## Abstract

**Background:** Bacterial cellulose (BC), natively synthesized by *Komagataeibacter* spp., is a biodegradable biomaterial with superior mechanical properties. However, under agitated cultivation, cellulose producing strain (Cel ^+^) often transition to non-producing mutants (Cel ⁻), restricting scalability and hindering widespread use. Agitation-associated shear stress, elevated oxygen levels, and genetic mutations have been linked to the emergence of the Cel⁻ phenotype. A genome-wide investigation that considers population heterogeneity and dynamics is essential to reveal the mutational landscape and evolutionary processes driving this phenotypic shift.

**Results:** Over successive rounds of agitated cultivation, the strain transitioned to a planktonic state, losing BC production. Whole-genome sequencing revealed both structural variations (SVs) and non-structural variations (NSVs). Contrary to previous reports, SVs, including insertion sequences (IS) mediated junction events, did not affect genes related to BC synthesis. Instead, the accumulation and positive selection of NSVs, such as frameshift and replication slippage events, in key BC-related genes, strongly correlated with the loss of cellulose synthesis

**Conclusions:** This study provides the first genome-wide population-level analysis revealing mutational dynamics underlying the BC phenotypic switch in *Komagataeibacter* spp. under agitated conditions. We show that BC-related gene mutations are not solely driven by SVs, with NSVs emerging as equally critical contributors. Furthermore, the genomic evidence implicates broader involvement of quorum sensing and cyclic dimeric guanosine monophosphate signaling, suggesting both genetic and regulatory factors underlying the disruption of BC synthesis. These findings highlight the importance of examining genetic heterogeneity with population to understand phenotypic adaptation for strain improvement strategies for scalable BC production.

## 1. Background

Bacterial cellulose (BC), primarily produced by *Komagataeibacter* spp. (formerly *Acetobacter* and *Gluconacetobacter*), has attracted growing interest in wide-ranging applications. Unlike plant-derived cellulose, high purity (devoid of lignin, hemicellulose, and pectin), superior physical and mechanical properties, along with the non-toxic and biodegradable nature has established BC as a sustainable biomaterial. (1)

The BC synthesis machinery, encoded by the bacterial cellulose synthase (bcs) operon, comprises of the core genes *bcsABCD*, flanked by accessory genes including *bcsZ* (endoglucanase, also referred to as *CMCax*)*, ccpAx* (cellulose complementing factor A), and *bglX* (beta-glucosidase) (2). Briefly, uridine diphosphate glucose (UDP-glucose), synthesized through the catalytic activities of phosphoglucomutase (*pgm*, converting glucose-6-phosphate to glucose-1-phosphate) and UTP-glucose-1-phosphate uridylyltransferase (*galU* converting glucose-1-phosphate to UDP-glucose), serves as the substrate for cyclic dimeric guanosine monophosphate (c-di-GMP) activated bcs operon that polymerizes the glucosyl residues to β- 1,4-glucan chains (2,3). Although *Komagataeibacter* spp. typically harbor multiple *bcs* operons (∼4-5 genome copies), only one constitutes a complete operon. A second, containing *bcsABC*, *bcsX* (putative SGNH/GDSL hydrolase), *bcsY* (putative acyltransferase), has been associated with the synthesis of amorphous or acylated cellulose, while the remaining operons generally encode only *bcsAB* or *bcsABC* genes. (4,5).

Despite the advancements in *Komagataeibacter* bioprocessing, scale-up of BC production remains constrained by the spontaneous emergence of non-cellulose producing (Cel^−^) mutants under agitated cultivation conditions (6). Conventionally, *Komagataeibacter* spp. are cultivated in static conditions. Being an obligate aerobe, the cells migrate along the dissolved oxygen gradient towards air-liquid interface producing BC pellicles (7). Prolonged synthesis at this interface hinders O_2_ diffusion, leading to a hypoxic environment (8) for the cells confined within the BC pellicle. To overcome this limitation, conventional shake flasks and bioreactors have been employed to enhance O_2_ mass transfer, for improved productivity (9). However, agitated conditions often generates random Cel^−^ mutants reducing BC production (10–12). Although the molecular basis remains unclear, shear stress and elevated oxygen in agitated cultures have been proposed as possible drivers of this phenotypic switch (6,10,12). One hypothesis is that unlike in static conditions where the pellicle facilitates O_2_ availability for cell growth at the air–liquid interface, BC synthesis during agitated cultivation provides no selective advantage. (10). Although limited, studies in *Komagataeibacter* spp. show that elevated O_2_ levels reduces *pgm* and *galU* (also referred as *gtaB*) (13) expression and increases phosphodiesterase activity, thereby decreasing the cellular c-di-GMP levels (14), a central regulator for motility, stress responses, virulence and biofilm formation (15).

Appearance of Cel^−^ mutants under shaking conditions has also been linked to both insertion sequences (IS)-mediated SVs and NSVs disrupting the BC-related genes (10). Coucheron et al. (16) reported an IS1031 disrupting the bcs operon of *K. hansenii* 23769 under agitated cultivation, while Hur et al. (17) observed a similar IS1031 insertion within the *bcsA* gene of *K. xylinus* DSM2325 under the same conditions. Furthermore, Krystynowicz et al. (12) demonstrated that in Cel⁻ *A. xylinum* E25 a single-base (T) deletion impaired both pgm and galU activity, thereby UDP-glucose synthesis.

Prior insights largely derive from Cel⁻ isolates (12,16,17), i.e., homogeneous populations that overlook the natural heterogeneity present under cultivation. This study applies the first population-level sequencing approach in *Komagataeibacter* spp. to dissect the mutational landscape of BC disruption under agitation. By mapping mutation hotspots and frequencies across heterogeneous populations, the study uncovers previously overlooked genetic and regulatory drivers of the Cel^−^ transition. The approach provides mechanistic insights into this phenotypic switch and actionable path for improving stability in BC production under agitated conditions.

## 2 Results

### 2.1 BC production during agitated cultivation

*K. intermedius* ENS15 (K.ENS15) was cultivated under agitated conditions in three parallel lines (Flasks A, B and C) for 10 successive Rounds of 3–5 days each (17) (see Methods). After the first Round, BC production was evident as a single agglomerated pellicle and the culture medium remained clear with no noticeable planktonic growth (Additional file 1: Fig.1). By the second Round, BC production persisted in all cultivation lines but was accompanied by visible planktonic growth. From this point onwards, BC production progressively declined, and by Round 6 it was completely absent in all lineages, with cells continuing to grow planktonically until the end of the experiment (Additional file 1: Fig.1). To evaluate cell growth, optical density at 600 nm (OD_600nm_) was measured at the end of each Rounds from both lysed BC pellicles and planktonic cultures. In Flask A OD_600nm_ values ranged between 1.1 and 3.3, corresponding to an average of 4.7 generations per Round. Flask B showed OD_600nm_ values between 1.1 and 2.6, averaging 4.5 generations per Round. Similarly, Flask C exhibited OD_600nm_ values between 1.1 and 2.6, with an average of 4.7 generations per Round (Additional file 1: Fig. 2).

### 2.2 Static cultivation to assess Cel^−^ to Cel^+^ reversion

Ability of Cel^−^ mutants to revert to Cel^+^ phenotype has been reported in literature (10). To assess the reversion capacity, cells from each agitated cultivation (either obtained after BC lysis or planktonic) were re-cultivated in static conditions, alongside the Cel^+^ parental strain. In contrast to the Cel⁺ parental strain (0.9 g/L BC), agitated cultures showed overall lower, fluctuating titres with occasional spikes: Flask A ranged from 1.1 g/L to 0.1 g/L, Flask B from 1.3 g/L to 0.1 g/L and Flask C from 1.3 to 0.3 g/L. (Additional file 1: Fig.3)

### 2.3 Population-level detection of insertion sequence-mediated SVs and junction events

SVs mediated by ISs and associated deletions have previously been documented in *Komagataeibacter* cells grown under agitated conditions (5,16–18). To investigate such events in K.ENS15 cultivations, the sequencing reads were initially aligned to the reference genome using the map-to-reference function in Geneious Prime 2025.0.3 (19), which highlights mismatches. However, this approach lacks sensitivity for detecting SVs in heterogeneous populations and cannot directly identify IS activity. Since the commonly used ISsaga (20) tool was unavailable, a combination of Breseq and ISEScan was employed to enable comprehensive detection of SVs and accurate annotation of IS elements (21–23). Prior analysis with the K.ENS15 reference genome (NCBI accession number, GCA_021555195.1) identified 73 IS elements. Figure 2 presents the IS identified in K.ENS15 by ISEScan, alongside those found in other characterized *Komagataeibacter* spp. A complete list of IS elements in K.ENS15, including genomic locations, is available in Additional file 2.

**Figure 2.**
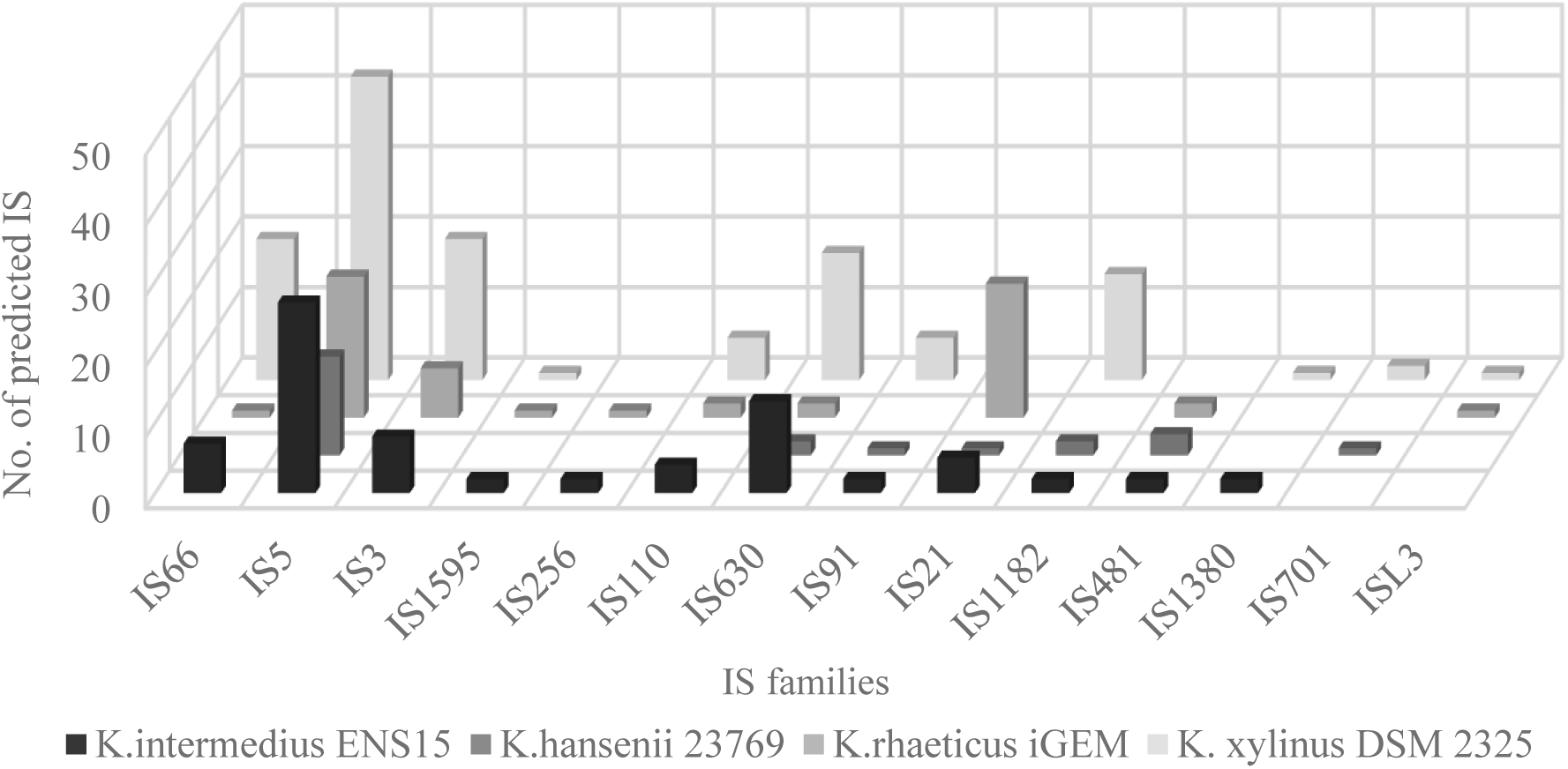
ISEScan predicted IS families distribution in *K. intermedius* ENS15, *K.hansenii* 23769, *K.Rhaeticus* iGEM and *K.xylinus* DSM2325

The unprocessed Breseq outputs are provided in Additional file 3, along with manually curated tables comparing the junction (JC) events between the parental strain and cells from the three lineages (Additional file 4). Majority of SVs detected by Breseq were categorized as “unassigned junction events”. JC events represent instances where two genomic regions, typically distant in the reference genome, appear adjacent in sequencing reads. These events frequently arise from IS activity leading to insertions, deletions, or duplications. The ‘unassigned’ designation indicates cases where the junction cannot be precisely resolved, often because it spans regions exceeding the sequencing read length or involves repetitive sequences.

Analysis revealed that most of the JC events were inherited from the Cel^+^ parental strain and persisted across the agitated cultivation lineages (Additional file 4). However, manual curation identified several JC events unique to the lineages, involving an IS element and a non-IS genomic region (Additional file 4). One example is a unique JC event detected after Round 10 in Flask C (Frequency 5.7%, Coverage 0.055), involving the IS1182_104 element (genomic position 2974235-2975693 bp) and an intergenic region of a gene encoding putative signalling protein and an oxygen-sensing gene (*dosp3*) (genomic position 936726-936776 bp) as shown in Figure 3. BLAST (24) analysis of the putative signalling protein identified it as a putative diguanylate cyclase/phosphodiesterase (DGC/PDE) domain-containing protein involved in cyclic-di-GMP signalling.

**Figure 3.**
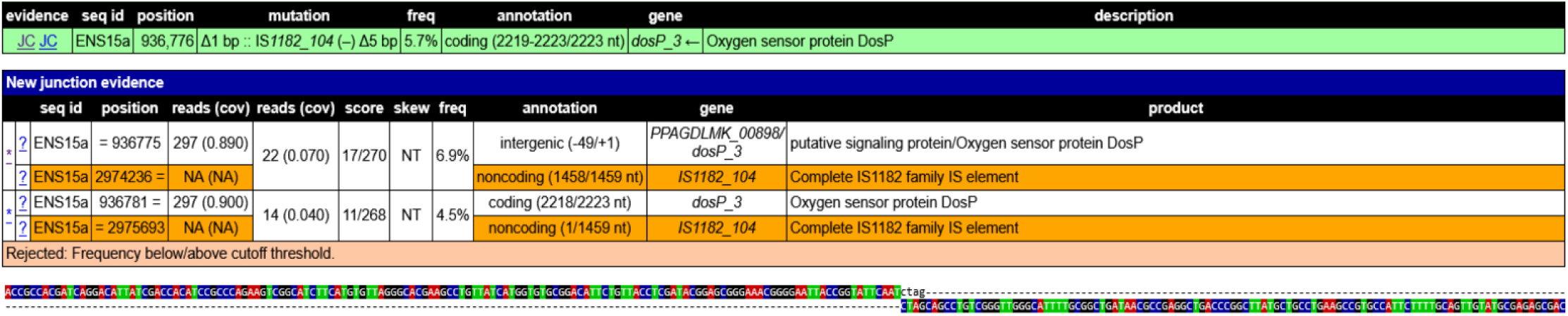
Breseq evidence of a JC event between IS1182_104 and the intergenic region between a putative signaling protein and an oxygen sensor protein. Rows correspond to IS1182 are highlighted in orange. In figure labels: *Position* refers to the genomic position of the interested sequence, the first *reads (cov)* show the amount of reads and the relative coverage of the sequenced region, the second *reads (cov)* reports the number of reads and the coverage supporting the mutation, and *Freq* denotes the percentage of reads containing the variant.

Previous studies have reported IS-mediated mutations to affect BC-related genes during agitated cultivation (16,17). Such loci was specifically examined in our dataset. The only IS-related event directly involving a BC gene was already present in the Cel^+^ parental strain and was inherited through the shaken lineages (Additional files 3 and 4). This event, supported by two JC events, was observed in the intergenic region between the *bcsA* (in bcs operon IV, genomic position 3299830-3304158 bp) (25) and an upstream putative acetyl-CoA synthase (genomic position 3304224-3306005 bp) (Additional File 1: Fig. 4 and 5). In the parental strain the two JC events were present with frequencies of 14.1% and 28.7 %. In the shaken lineages, these frequencies fluctuated between 11.5–21.2% and 23.4–35.9% in Flask A; 10.0–19.1% and 22.5–36.4% in Flask B; and 12.2–22.6% and 24.3–36.2% in Flask C (Additional files 3 and 4).

### 2.4 Population-level detection of NSVs

Consistent with the JC events observations, most NSVs were inherited from Cel^+^ parental strain, although mutations unique to the shaken lineages were also identified. A comprehensive comparison of all NSVs detected in both the parental strain and shaken cultures, along with non-“unassigned” JC events are presented in Additional file 5. Most NSVs unique to the agitated cultivations showed no evidence of positive selection over time, instead they appeared after specific cultivation rounds and were not retained long-term. For instance, in Flask A and C, SNPs were detected in *ydaD_1* encoding general stress protein 39 (genomic position 2297035-2297895 bp). In Flask A, after Round 1, a G→A substitution at genomic position 2297613 was observed at 10.4% frequency, causing an R95W amino acid change. A similar SNP appeared in Flask C after Round 7 at 9.3% frequency. Another G→A substitution at genomic position 2297622 bp, detected in Flask A after Round 8, was synonymous and did not cause an amino acid change. However, a G→A change resulting in an R90C amino acid substitution was identified in Flask C after Round 7 at a frequency of 7.2%.

#### 2.4.1 NSV events within key genes involved in BC synthesis

As anticipated based on prior studies, NSV events were detected in several BC-related key genes, including *ccpA, bcsAI, bcsCI, galU,* and *pgm* (Table 1). Several of these mutations, absent in Cel^+^ parental cells, displayed evidence of positive selection over time, with some recurring independently across all three lineages.

**Table 1.**
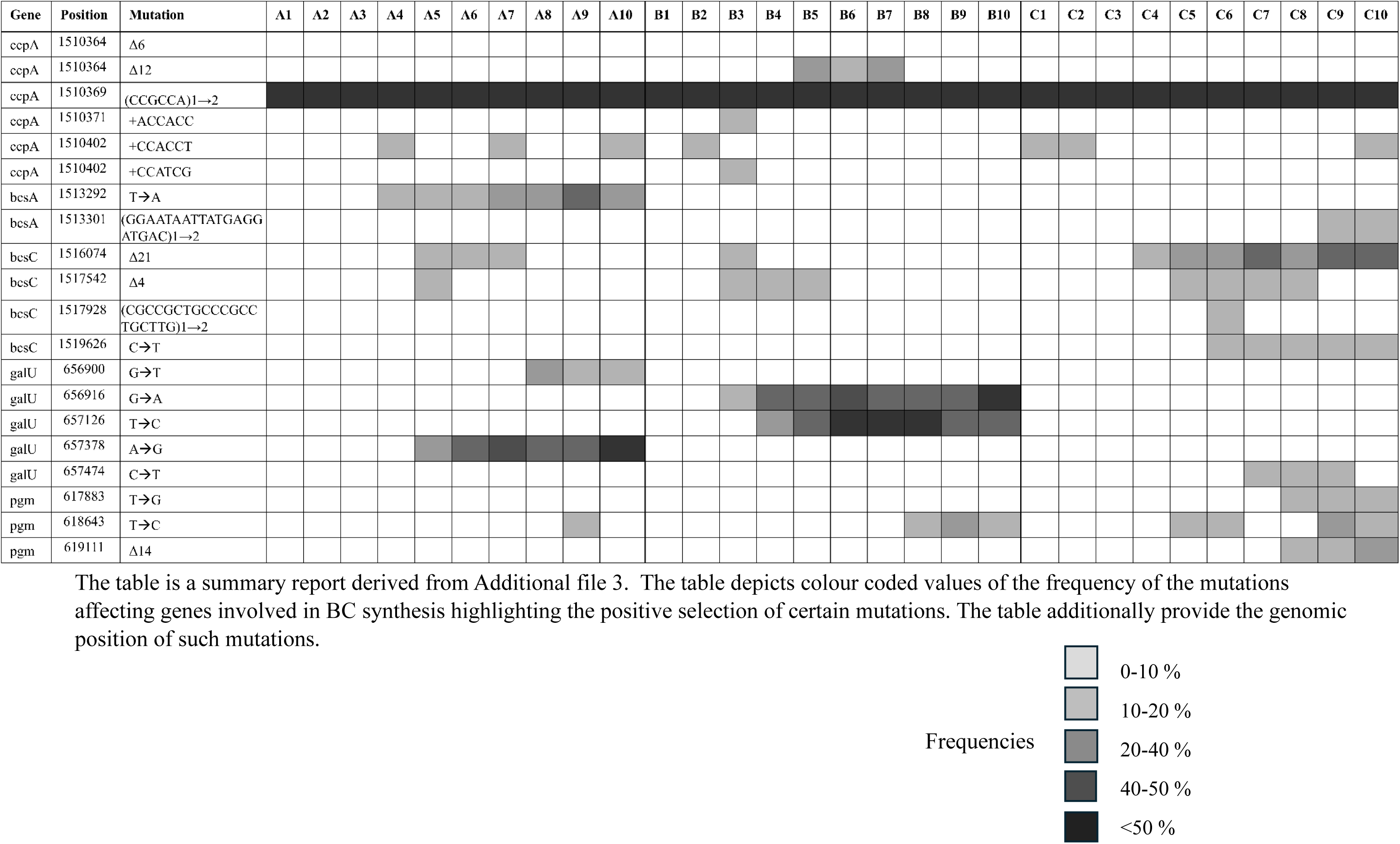
NSV events within key genes involved in BC-synthesis.

Notably, most NSVs were clustered within the accessory gene *ccpA*, specifically within a hotspot region (genomic position 1510364–1510402 bp) encoding 13 consecutive proline residues derived from repeated CCG/CCA triplets prone to replication slippage (Additional file 1:Fig.6) (26). A parental slippage event, expanding one to two copies of the CCGCCA repeat, was retained at high frequency (83–88%) across all lineages (Table 1). Additional lineage-specific insertions also arose within this hotspot. In Flask A, a +CCACCT insertion (adding two prolines) appeared after Rounds 4, 7, and 10 with frequencies ranging from 5.6% - 7.8%. In Flask B, the same insertion was detected after Round 2 (8.8% frequency), alongside two further insertions, +ACCACC (5.1% frequency) and +CCATCG (5.9% frequency), introducing two threonine and a proline-serine pair, respectively. Flask C showed a heterogeneous population carrying the +CCACCT insertion at Rounds 1 and 2 (with 5.8% and 5.0% frequencies, respectively), disappearing in intermediate rounds, and re-emerging at Round 10 (8.9% frequency). Additionally, small deletion of 6–12 bp removing 2 – 4 proline residues (genomic position 1510364-1510369/1510375 bp) appeared transiently in Flasks B and C at frequencies between 12–18% during the early rounds but were not positively selected by Round 10.

Within the *bcs* operon, NSVs were detected exclusively in *bcsA* and *bcsC*. In *bcsA*, a nonsense mutation (TAT → TAA) introducing a premature stop codon (genomic position 1513292 bp) emerged in Flask A. This mutation, absent in Flasks B and C, was gradually enriched over the course of cultivation, increasing in frequency from 9.4% at Round 4 to 15–36% by Rounds 9 and 10. In Flask C, a 20 bp duplication (GGAATAATTATGAGGATGAC, genomic position 1513282-301 bp) in *bcsA,* causing a frameshift and emergence of two stop codons, was detected in early rounds but was lost by Round 10 (Additional File 1:Fig.7; Table 1). Overall, *bcsA* variants were observed only in Flasks A and C, with Flask A’s stop codon mutation predicted to disrupt the entire BC synthase complex, while Flask C’s frameshift mutation as transient.

In contrast to *bcsA*, the *bcsC* gene was affected across all lineages. Two recurrent mutations were identified: a 21 bp deletion (GCACGTTTCTGGTTGCAGCAG, genomic position 1516074-94 bp) and a 4 bp deletion (GGCA, genomic position 1517542-45 bp). The 21 bp deletion removed the amino acid sequence ARFWLQQ within a tetratricopeptide repeat (TPR) region of *bcsC*. Using the freely available SwissModel tool (27), we created a 3D model of the mutated *bcsC*, to compare it against the wild type *bcsC* (Additional File 1:Fig.8). The 4bp deletion caused a frameshift and introduced premature stop codons (Additional File 1:Fig.9). Although the timing of these deletions varied across lineages, the 21 bp deletion was positively selected only in Flask C, with population-level frequencies increasing from 5.7% (Round 4) to 35% (Round 10). In contrast, the 4 bp deletion remained at low frequencies (6–10%) and showed no evidence of positive selection (Table 1). Additional events were also identified in Flask C. A duplication event (CGCCGCTGCCCGCCTGCTTG_1_→_2_, genomic position 1517909-928 bp) after Round 6 causing a frameshift and multiple stop codons but was not retained in subsequent rounds (Additional file 1: Fig.10). In the same round, a SNP (CAG→TAG, genomic position 1519626 bp) introduced a premature stop codon, which persisted at low frequencies (6% – 9%) within the heterogeneous population until Round 10.

NSV events were identified in *galU*, encoding UTP–glucose-1-phosphate uridylyltransferase, essential for UDP-glucose formation and BC synthesis. These mutations appeared across all shaken lineages, with varying type and temporal dynamics (Table 1). For instance, in Flask A, an A→G substitution resulting in a tyrosine-to-cysteine amino acid change at position 172 (Y172C) emerged at Round 5 (frequency, 19%) and was positively selected to 75% by Round 10. Although a second mutation causing an alanine-to-serine substitution at position 13 (A13S) appeared, this was detected to be transient and was not selected. Flask B showed two additional point mutations absent in Flask A: a G→A substitution causing arginine-to-histidine to change at position 18 (R18H), which was detected at Round 3 with 8% frequency was positively selected by Round 10 (frequency, 72%), and a T→C substitution causing leucine-to-proline to change at position 88 (L88P), reaching a frequency of 63% at Round 7 before declining and stabilizing at 23% by Round 10. Although similar mutations were absent from heterogenous populations in Flask C, a nonsynonymous SNP (S204F) was detected between Rounds 7 and 9, which was absent by Round 10.

NSV events were identified in the *pgm* gene, which encodes phosphoglucomutase, the enzyme catalyzing the conversion of glucose-6-phosphate to glucose-1-phosphate, a key step in UDP-glucose biosynthesis. Mutations in *pgm* were observed across all shaken lineages with distinct frequencies and patterns (Table 1). Three key NSV events were detected within *pgm*: two point mutations and one deletion. A T→C substitution causing a lysine-to-glutamate amino acid change at position 158 (K158E), was present in Flasks B and C at low frequencies (5% – 16%), but absent in Flask A. The other two SNV variants, a T→G mutation resulting in an aspartate-to-alanine change at position 411 (D411A) and a 14bp deletion (GCATTGACTGCTCC, bp 619111-124) removing the start codon, were identified exclusively in Flask C from Round 8 onwards (Additional File 1: Fig. 11). The deletion and mutation frequencies were initially 10.4% and 8.4%, respectively and they were retained until the final cultivation round with frequencies of 9.2% and 18.5% (Additional File 5: Table 2).

## 3 Discussion

Our empirical observations during shaking cultivations confirmed a progressive reduction in BC production, accompanied by increased planktonic cell growth, consistent with previous studies (28,29). This phenomenon has been linked to mutations identified in homogenous population isolated as Cel^−^ mutants on agar plates, (12,16,17,30). For example, Krystynowicz et al.(12) reported a SNP causing reduced expression of *pgm* and *galU* (also referred to as *gtaB*) following shaking cultivation of *K. xylinus* E25. Similarly, Coucheron et al. and Hur et al.(16,17) reported IS1031 element insertions disrupting the *bcs* operon in different *K. xylinus* strains after agitated cultivations.

Unlike approaches that isolate single colonies and detect mutations at 100% frequency, our study uniquely analysed mutations from the perspective of a natural heterogenous cell population, providing deeper insights into the mutation dynamics under agitated conditions. This population-level approach preserves the broader genetic diversity context. We observed that BC synthesis genes were affected across all three lineages, with some mutations undergoing clear positive selection over time, indicating potential growth advantages. Notably, no mutations were detected within genes involved in core metabolic pathways.

Contrary to the prior find by Coucheron et al. and Hur et al. (16,17), we found no evidence of IS elements disrupting BC production in our experiments. However, an IS element affecting *bcs* operon IV was identified in the Cel^+^ parental strain, which retained BC-producing capability (Additional file 1: Fig. 5 and 6). Comparative analyses suggest this operon resembles those implicated in amorphous cellulose synthesis (5).

NSVs in key BC synthesis genes (*pgm*, *galU*, and *bcs* complex) were consistent with previous findings (Table 1) (31). Earlier studies in Cel^+^ *K. oboediens* demonstrated that deletions of 5 and 17 bp could generate Cel⁻ phenotypes, while repeated static cultivation restored BC production via compensatory 2 bp insertions that nullified the frameshift (31). Our population-based approach revealed that the mutations associated with impaired or absent BC production were present at low frequencies or not detected. Furthermore, each flask exhibited distinct mutational landscapes, suggesting additional mechanisms uniformly suppress BC production independent of predominant genotypes.

Recently, Zhang et al.(32) reported *K. xylinus* 2955 to possess a complex c-di-GMP regulatory mechanism, comprising of 10 genes with tandem PDE/DGC activity (GGDEF−EAL domain proteins). The study established a direct linkage between the regulatory c-di-GMP levels with BC production, reporting that the knockout of *pde*, which regulates the degradation of total cellular c-di-GMP, led to a 48% increment in BC synthesis. Complementary findings by Tuckerman et al.(14) in *E. coli* demonstrated a positive correlation of elevated O_2_ levels to enhanced PDE activity, accelerating c-di-GMP degradation (14). Similarly, Huang et al. reported that increased O_2_ concentrations led to a downregulation of the BC pathway gene expression in *K. xylinus* CGMCC 2955 (13,14).

Our BLAST analysis identified nine loci encoding putative PDE/DGC activity in K.ENS15. Cannazza et al. (25), originally annotated seven of these as oxygen-sensing sensor proteins (*dosP*), indicating active modulation of c-di-GMP metabolism in response to O_2_. (Additional file 1: Table 1). No mutations affecting these genes were observed within the shaken lineages, except a late-round mutation in *dosP_3* in Flask C, which appeared too late to influence BC production (see section 2.3).

Based on this mutational landscape and the empirical data, we propose c-di-GMP regulation as a key mechanism driving BC loss in agitated cultivations. Elevated O_2_ levels likely enhance PDE activity, lowering the total c-di-GMP levels and favouring the selection of genotypes with BC production due to their growth advantage (6,12). This hypothesis is supported by the reversible nature of Cel⁻ to Cel⁺ phenotypic switch observed under static cultivation, correlating to the changes in O_2_ availability (29). To evaluate this hypothesis, we performed a preliminary experiment to quantify the total intracellular c-di-GMP levels in cells from Round 1 and Round 10 of Flask A lineage (33). Preliminary fluorometric measurements indicated a 28% ± 4% decrease in total intracellular c-di-GMP (Additional File 1: Fig.12). While this supports the hypothesis, global c-di-GMP levels may not fully reflect the localized pools critical for BC regulation (32). Future work will include transcriptomic and metabolic evaluations to elucidate c-di-GMP signalling dynamics under agitated conditions.

Furthermore, quorum sensing (QS) mechanisms may also influence BC regulation during agitated cultivations. N-acyl homoserine lactone (AHL) QS has been documented in *Komagataeibacter* spp. (34,35). K.ENS15 contains a putative *LuxR* homologue (genomic position 2014782-2105501 bp), but lacks the canonical *LuxI* synthase, consistent with other *Komagataeibacter* spp. (36). Despite this, QS regulation of PDE activity via alternative signalling routes has been reported in *Komagataeibacter spp.* (35). Additionally, Valera et al.(37) demonstrated quorum quenching (QQ) activity in *K. europaeus* via the AHL-degrading GqqA protein (also present in K.ENS15, genomic position 635676 - 636521 bp), which when supplemented BC production (37). These studies highlights the significant roles of QS and QQ in modulating BC synthesis and understanding these regulatory pathways is vital in improving the process scalability.

Despite meticulous handling; static cultivation for BC synthesis, minimal-agitation BC lysis, and cryopreservation for glycerol stock preparation; the presence of IS insertions and mutations in the parental strain indicates genomic divergence from the originally isolated strain. This underscores the inherent genomic instability of *Komagataeibacter* spp. and emphasizes the need for rigorous sequence verifications when working with these bacteria. Future studies will extend to additional strains and glycerol stocks, employing long-read sequencing approaches to improve the resolution of IS events.

## 4 Conclusions

This study presents the first comprehensive characterization of the mutational landscape within a heterogeneous population of *Komagataeibacter* spp. undergoing the well-documented Cel^−^ to Cel^+^ phenotypic switch under agitated cultivation. Our results indicate that the loss is driven not primarily by mutations in BC biosynthetic genes but by regulatory mechanisms, with the c-di-GMP signalling pathway playing a central role in O_2_ -responsive fluctuations caused by agitated cultivation. However, more detailed studies are required to quantify c-di-GMP dynamics, regulatory gene expression, and the contributions of QS and QQ mechanisms on BC synthesis. By analysing heterogeneous populations instead of isolated clones, our approach reveals population-level mutational dynamics of *K. intermedius*, and notable genomic divergence over time. Together, these findings highlight the importance of dynamic regulatory responses over structural mutations in controlling BC production during agitated cultivation conditions. Future transcriptomic and metabolic analyses are essential to clarify c-di-GMP dynamics and QS/QQ roles.

## 5 Materials and Methods

### 5.1 Chemicals and materials

GeneJet genomic DNA purification Kit was acquired from Fisher Scientific (Vantaa, Finland). Citric acid, glucose, sodium chloride and sodium hydroxide were purchased from VWR international Oy (Helsinki, Finland). Cellulase from *Trichoderma reesei,* disodium hydrogen phosphate, potassium dihydrogen phosphate and potassium chloride were acquired from Merck (Espoo, Finland). Yeast extract and bacteriological peptone were acquired from Neogen (Lansing, USA). The cyclic-di-GMP Assay kit was purchased from Lucerna (New York, USA). Black, flat bottomed 96-well plates were purchased by VWR international Oy (Helsinki, Finland).

### 5.2 Strain, media and agitated cultivation

K.ENS15 was isolated by Cannazza et al. (25). For preculture preparation, 100 µL of glycerol stock (parental strain) was inoculated in 10 mL of HS-glucose medium (g/L; 5 peptone, 5 yeast extract, 2.7 Na_2_HPO_4_, 1.15 citric acid and 20 g/L glucose) and incubated in static condition for 5 days at 30 °C. The synthesized BC was lysed overnight using 2 % cellulase to release the entrapped cells, which were subsequently washed thrice with sterile 1X Phosphate buffered saline (PBS, g/L, 8 NaCl, 0.2 KCl, 1.44 Na_2_HPO_4_, and 0.24 KH_2_PO_4_). (25)

The washed cells were used as inoculum for agitated cultivations (initial OD600 of 0.08), conducted in 250mL Erlenmeyer flasks, with three parallel cultivation lines designated as Flask A, Flask B and Flask C. Cultivations were conducted in 50 mL HS-glucose media at 30°C and 230rpm for 10 consecutive rounds, each lasting 3-5 days (17). At the end of each Round, 1 mL of cellulase was added irrespective of observable cellulose formation. The cultures were subsequently washed with PBS as previously described(25). The optical densities of the resulting heterogenous cell suspensions were measured and used both as inoculum for the following round of agitated and static cultivation and as a source for sequencing. A flask with containing HS/glucose media was included in each round as a contamination control.

### 5.3 Static cultivations and BC processing

Static cultivations were carried out in 50 mL culture tubes, with total 10 mL volume consisting of HS-glucose medium and inoculum. The static cultivations were seeded to start with an OD of 0.08. and the tubes were incubated for 4 days at 30°C. BC pellicles were processed as mentioned in Cannazza et al., 2021. (38)

### 5.4 Genome sequencing, variants calling and bioinformatics

Genomic DNA was extracted using GeneJet Genomic DNA Purification Kit as per manufacturer’s instructions (Fisher Scientific, Finland). Sequencing was performed at Novogene Gmbh (Planegg, Germany) using the Illumina NovaSeq X Plus system, generating approximately 1 Gb of raw data per sample. The reads quality was assessed using FastQC (v0.12.1) (39) and trimmed using Trimmomatic (v1.39) (40). Genetic variation analysis was conducted using Breseq (v0.39.0)(21,22,41–43) in polymorphism mode, with the deposited K.ENS15 reference sequence (GenBank accession: GCA_021555195.1). Prior to variant calling, insertion sequences in the reference genome were annotated using ISEScan (v.1.7.2.2) (23). This step proved to be essential since the widely used ISaga (44) was unavailable (accessed 2024 - 2025). Detailed instructions for integrating ISEScan with Breseq are described in the Breseq user manual (41). Genomic samples analysed included those collected at the end of each cultivation round as well as the parental strain. To compare the mutational landscapes, Breseq raw outputs were further processed using GDtools (41) and examined in manually curated tables generated with Python 3 scripts. Specifically, NSV and the JC events not classified as “unassigned” were compared using GDtools (41). In contrast, “unassigned junction events,” typically linked to IS activity, were manually curated by selecting cases connecting an annotated IS element with another genomic region. Events occurring exclusively between two non-annotated regions, or between a non-IS annotated region and an unannotated one, are provided in the raw Breseq outputs (Additional file 3).

The bioinformatics analyses were performed on the Puhti supercomputer provided by CSC(45). Geneious Prime 2025.0.3 (19) supported initial variant identification (see section 2.2), visualization of genomic regions of interest, and functional assessment of mutations. The Basic Local Alignment Search Tool (BLAST) provided by NCBI (24) was utilized to examine sequences of interest and manually annotate hypothetical proteins.

### 5.5 C-di-GMP fluorometric assay

The c-di-GMP fluorometric assay was conducted according to supplieŕs instructions, using black, flat bottomed 96-well plates with one modification. To enhance accessibility of intracellular c-di-GMP, cell pellets obtained from PBS-washed cultures at the end of the shaking cycles were briefly sonicated at 20% amplitude, after resuspension in 4 mL of 1X PBS (33). Fluorescence measurements were conducted using a Biotek Cytation 3 with emission at 485 nm and excitation at 528 nm. Background fluorescence was subtracted from recorded values and normalized by OD_600nm_ values of the resuspended pellets. A parallel negative control, in which all the reagents were added except for the c-di-GMP sensor molecule, confirmed that the detected fluorescence signal originated from intracellular c-di-GMP.

## Declarations

## Acknowledgments

The authors thank M.Sc. Efstathia Mantzari for valuable discussions on the bioinformatic setup. D.B and R.M acknowledges the Center for Young Synbio Scientists and the Research Council of Finland (Grant Number 346983) for supporting this study. D.B further acknowledges CSC – IT Center for Science, Finland, for providing the computational resources.

## Availability of data and materials

All the raw data, except the c-di-GMP data, are included in the additional files. The c-di-GMP raw data are available under request. Additional File 3 is available upon request, as its large data volume prevents uploading.

## Competing interest

The authors declare no competing interests.

## Funding

This work has been done at CYSS Center for Young Synbio Scientists with support from Jenny and Antti Wihuri foundation.

## Authors contribution

D.B. designed and performed the experiments, including shaking cultivations, bioinformatics, and genome analysis; interpreted the data; and drafted the original manuscript. R.M. conceptualized the project, interpreted the data, revised the manuscript, and supervised the work.

## Notes

### Competing Interest Statement

The authors have declared no competing interest.

