## Additional File 1 for "Exploring Genome-Wide Mutation Dynamics and Bacterial Cellulose Impairment in *Komagataeibacter intermedius* Cultivated Under Agitation Stress"

Davide Bersanetti 0009-0001-9987-4220

Rahul Mangayil 0000-0001-7547-409X

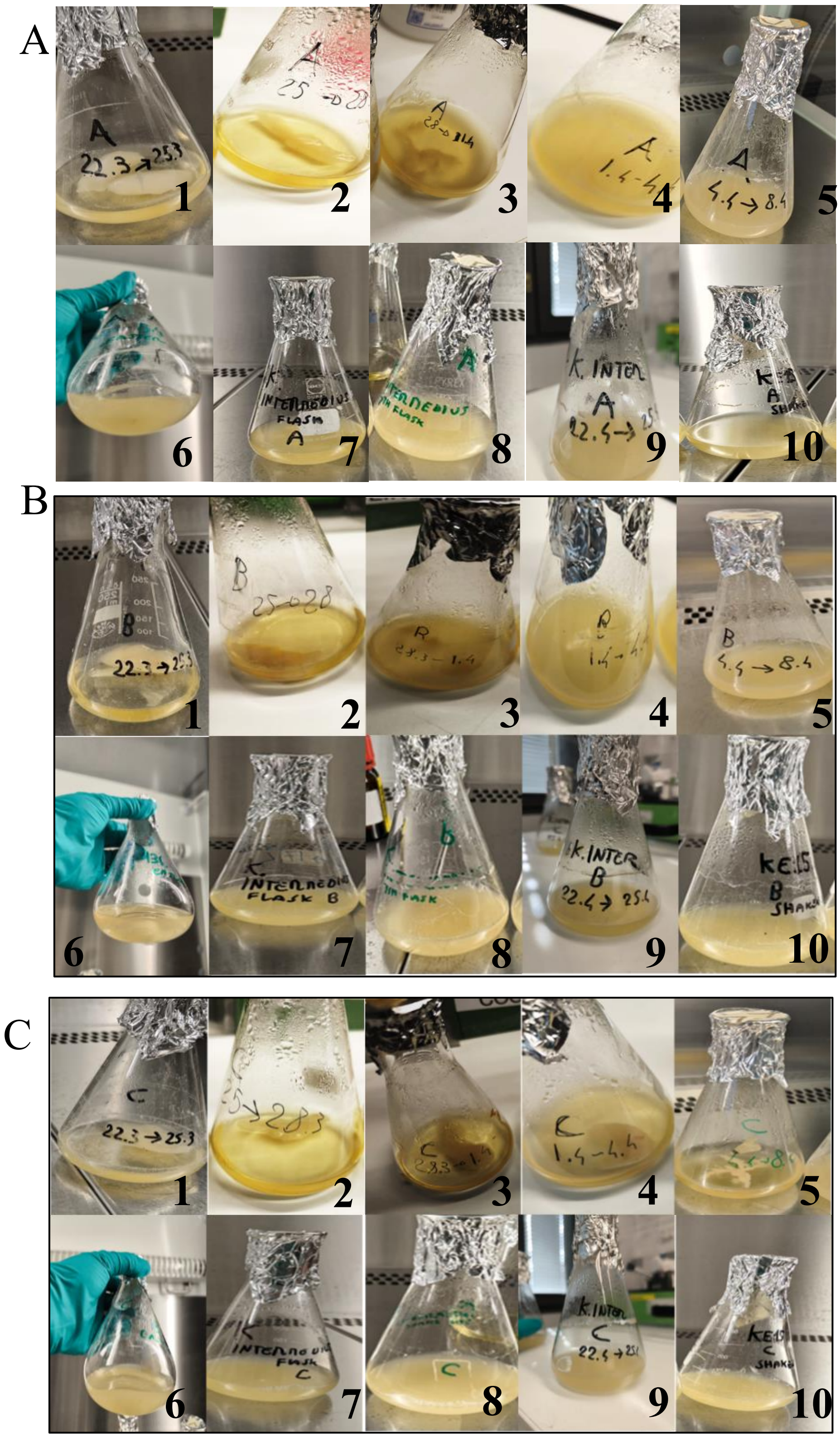


Figure 1: BC synthesis and planktonic growth of K. ENS15 lineages under agitated cultivation. A) Flask A over ten rounds (1–10), showing complete loss of cellulose production by Round 4. B) Flask B over ten rounds (1–10), similarly exhibiting no cellulose production starting at Round 4. C) Flask C over ten rounds (1–10), with cellulose clearly produced until round 5 but completely halted from Round 6 onward.

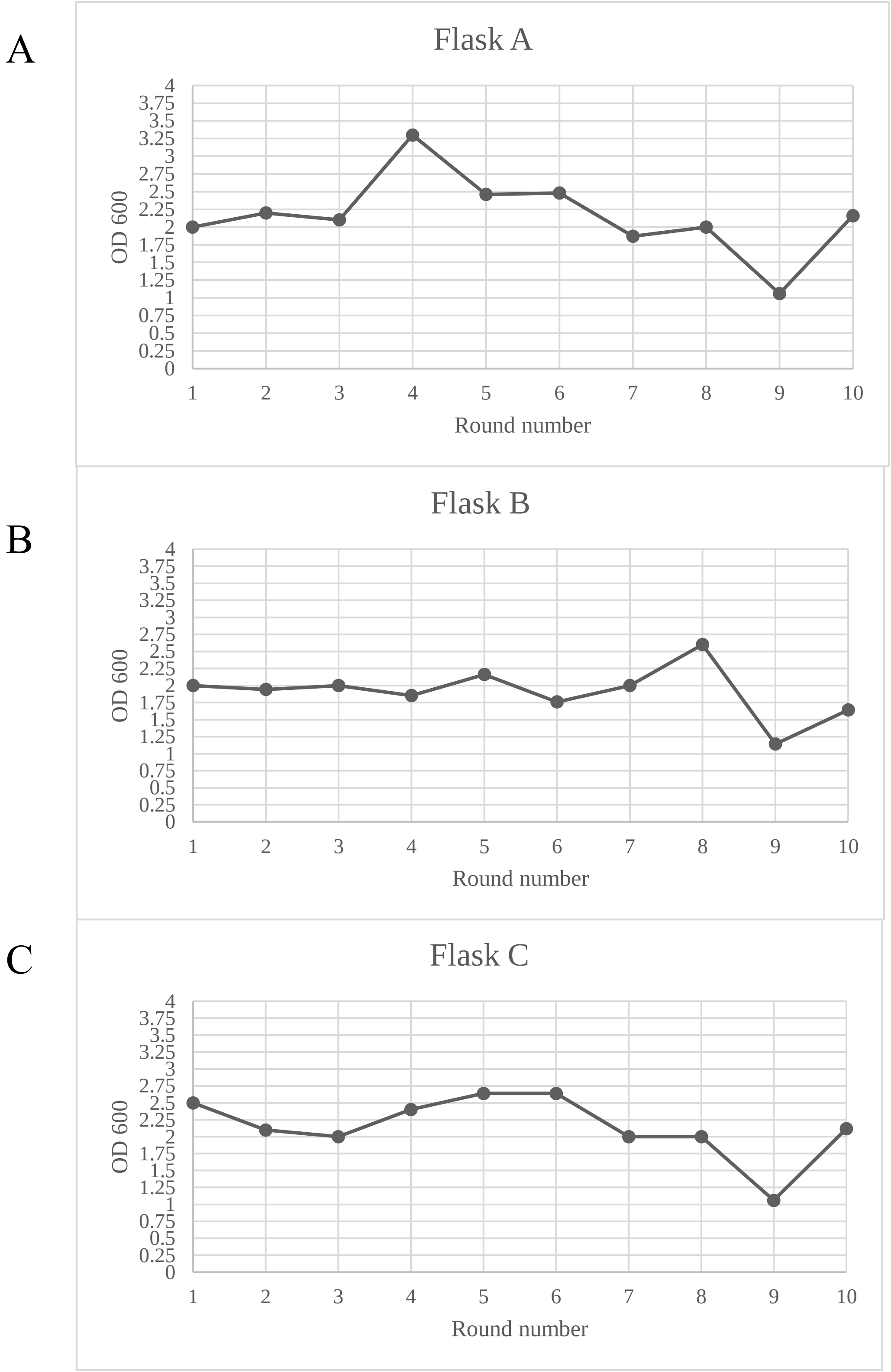


Figure 2. Cell growth ($OD_{600nm}$) measurement across 10 cultivation rounds for all three cultivation lines. A) represent the OD600nm measurement for Flask A. B) represent the OD600nm measurement for Flask B. C) represent the OD600nm measurement for Flask C

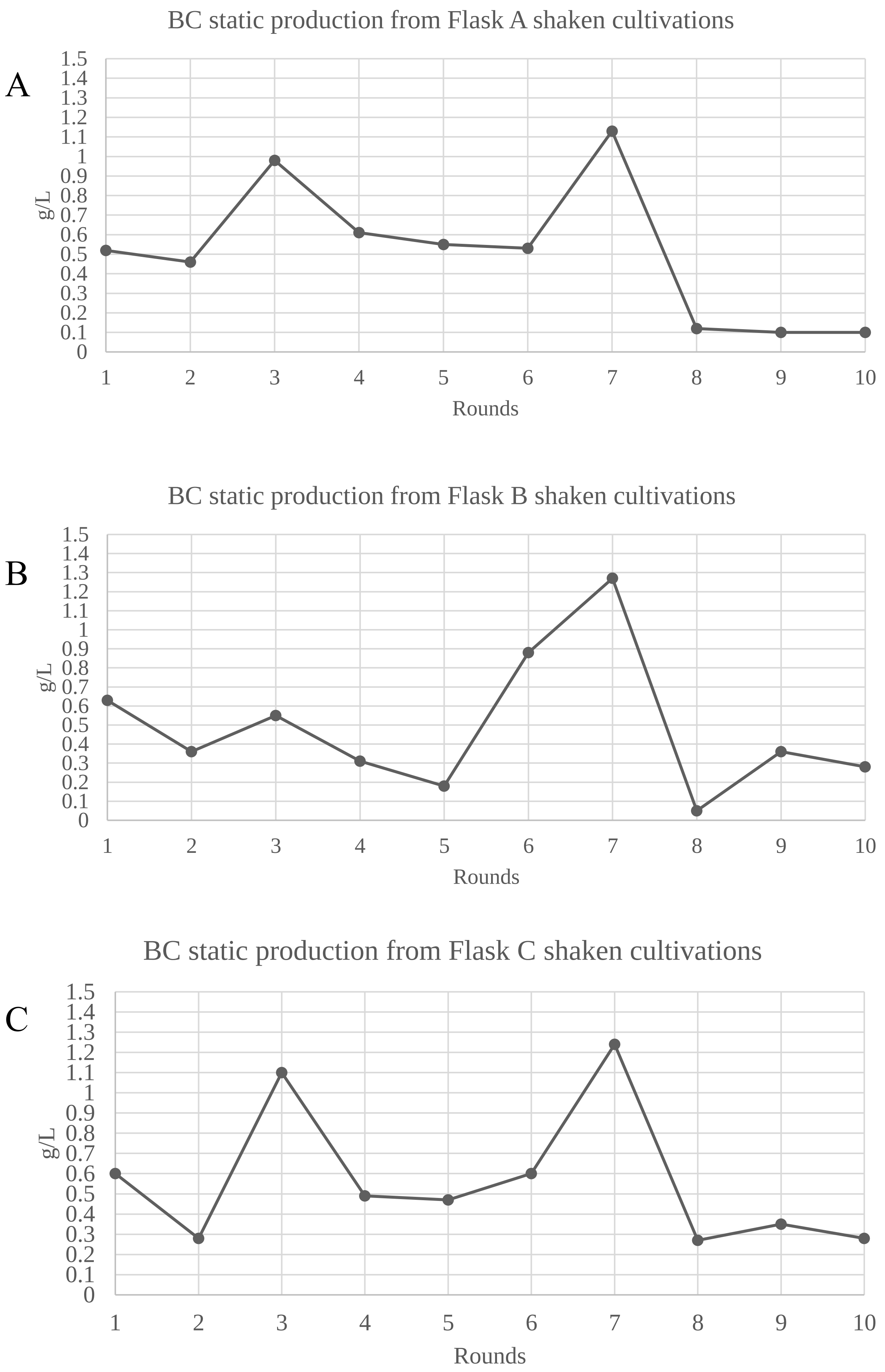


Figure 3. BC cellulose production ( reported as g/L) achieved by static cultivations inoculated with cells obtained from the shaken cultivations. A) production achieved in Flask A. B) production achieved in Flask B. C) production achieved in Flask C.

**New junction evidence**

| | seq id | position | reads (cov) | reads (cov) | score | skew | freq | annotation | gene | |
|---|---|---|---|---|---|---|---|---|---|---|
| ? | ENS15a | = 3090027 | 679 (1.800) | 55 (0.150) | 21/274 | NT | 14.1% | intergenic (+565/ +487) | *PPAGDLMK_02869/ xpkA* | hypothetical protein/Xylulose-5-phosphate phosphoketolase |
| ? | ENS15a | 3304182 = | 0 (0.000) | | | | | intergenic (-24/ +42) | *acsAB_3/acsA* | Cellulose synthase 1/Acetyl-coenzyme A synthetase |

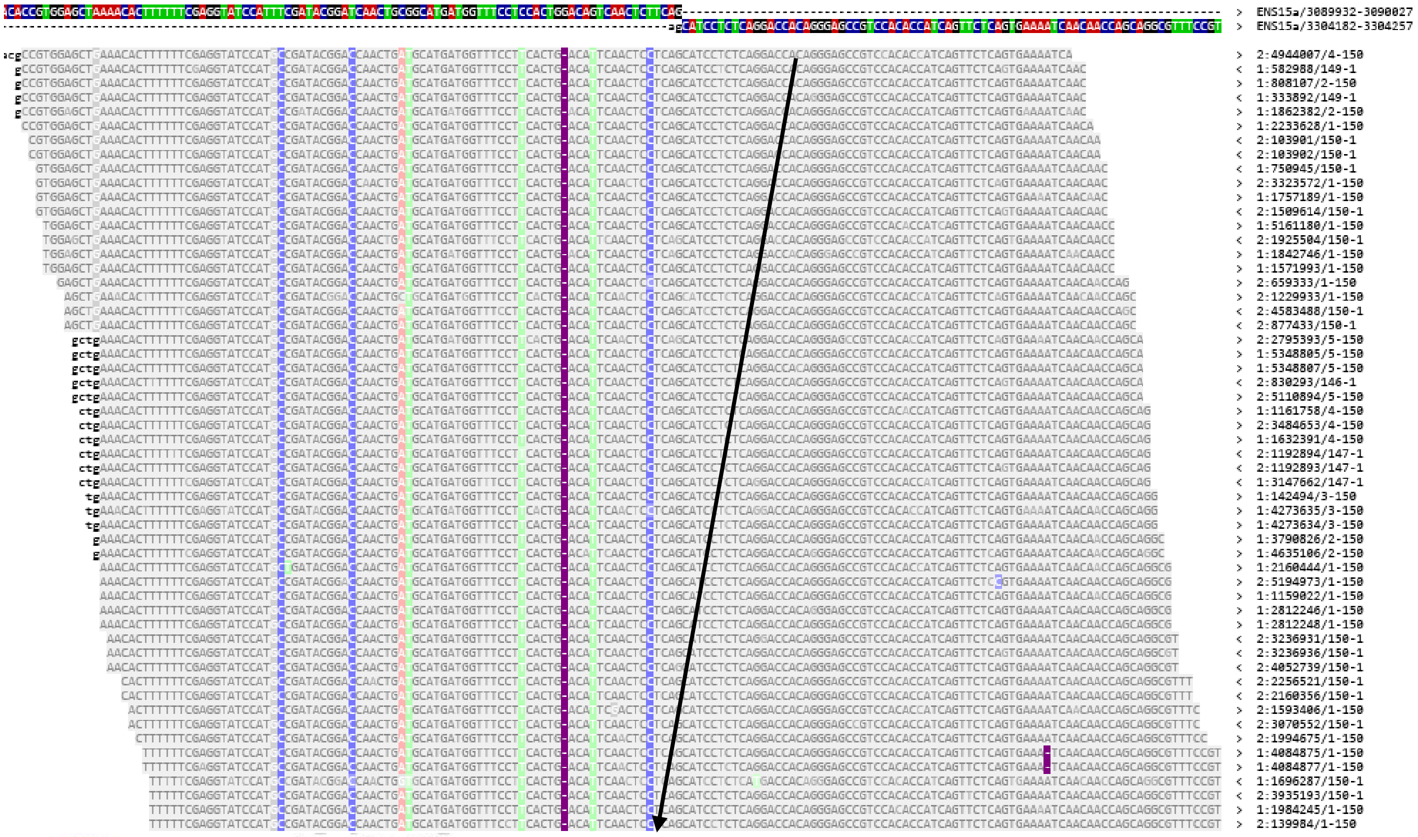


```
CDS             complement(<1..56)
                /locus_tag="SCD25_11865"
                /inference="COORDINATES: similar to AA
                sequence:RefSeq:WP_010336579.1"
                /note="Derived by automated computational analysis using
                gene prediction method: Protein Homology.
                GO_function: GO:0004803 - transposase activity [Evidence
                IEA]"
                /codon_start=1
                /transl_table=11
                /product="IS110 family transposase"
                /protein_id="WPP23542.1"
                /translation="MHQLVRIGMDTSKKVFQLHGVDAEERVALSRKLSRHQMIRFFEK
                LPPTVIGIEACGASHHWARTLGALGHEVRLMAPQLVKPYVRRSKNDAADAEALCEAMS
                RPTMRFVPVKTVEQQAALMLVGMRERLVGRRTQLANSIRGYAAEFGLTAPLGLSRVEP
                FLRQIASEDILPSLAREMFAQMAEEYGALMEHIRELDEKLATWHRSQEVSRRLSGIPG
                IGPLGAALLLMKAPDPHLFRSGRDFAAWIGLTPRDHSSGGKVRHGGITRAGDSRLRST
                LVVGATTVLRHVRQNKQSRLATPWLVSLLGRKKAKLAAVALANRIARIAWKLMTSGET
                YRKPVAVDAMTAATAS"
gene            143..>148
                /locus_tag="SCD25_11870"
                [illegible]
```

Figure 4. Reads supporting a JC affecting the bcs operon IV in the parental strain. These reads, when analyzed with BLAST, match a transposase sequence.

**New junction evidence**

| | | seq id | position | reads (cov) | reads (cov) | score | skew | freq | annotation | gene | |
|---|---|---|---|---|---|---|---|---|---|---|---|
| * | ? | ENS15a | 3304180 = | 0 (0.000) | 126 (0.440) | 51/212 | NT | 28.7% | intergenic (-22/ +44) | *acsAB_3/acsA* | Cellulose synthase 1/Acetyl-coenzyme A synthetase |
| | ? | ENS15a | 3495571 = | 625 (2.170) | | | | | intergenic (+55/ +126) | *PPAGDLMK_03263/ PPAGDLMK_03264* | hypothetical protein/IS256 family transposase ISSpwi2 |

Only 100 of 133 total aligned reads displayed.

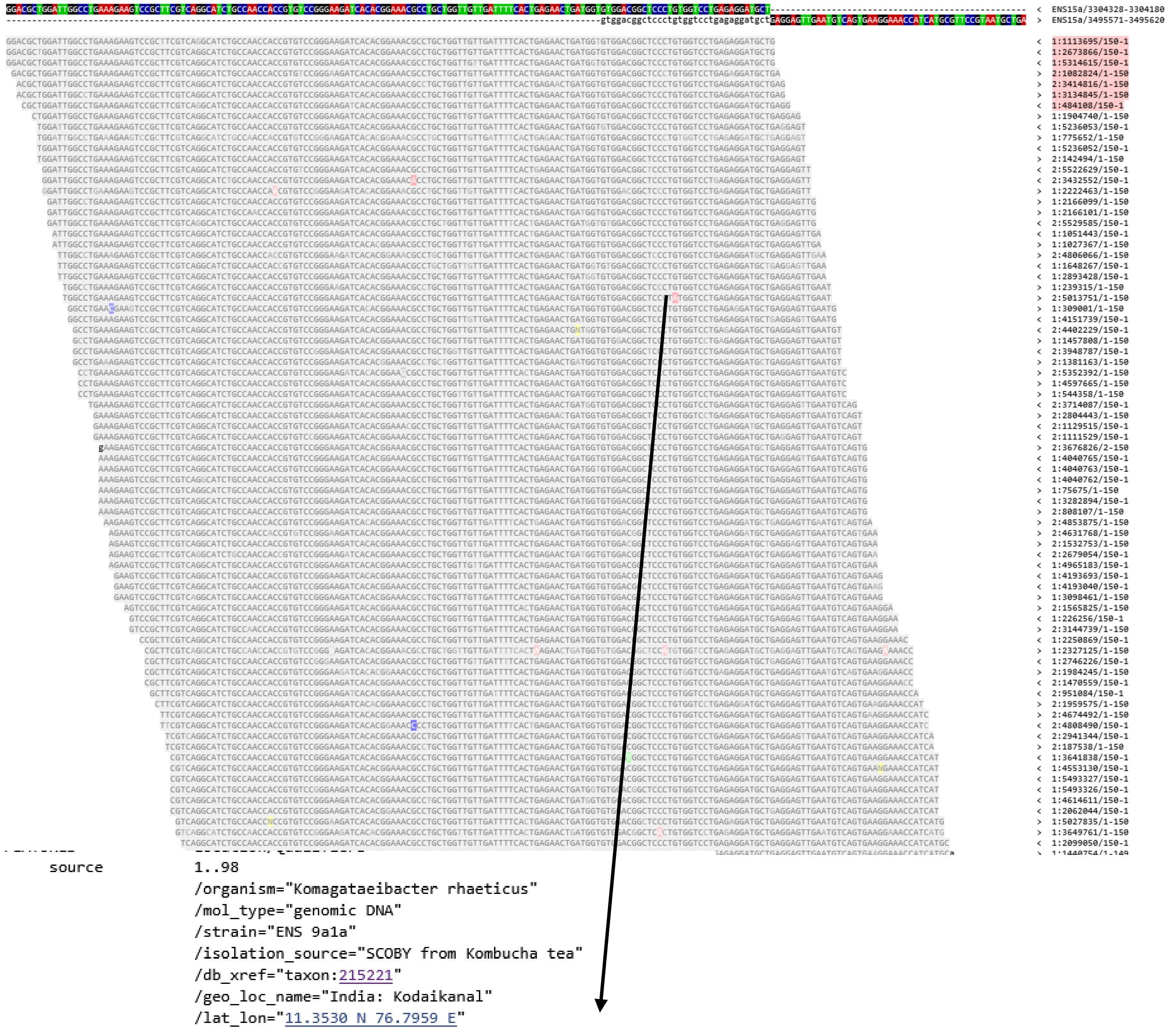

```
     source          1..98
                     /organism="Komagataeibacter rhaeticus"
                     /mol_type="genomic DNA"
                     /strain="ENS 9a1a"
                     /isolation_source="SCOBY from Kombucha tea"
                     /db_xref="taxon:215221"
                     /geo_loc_name="India: Kodaikanal"
                     /lat_lon="11.3530 N 76.7959 E"
                     /collection_date="Aug-2019"
     gene            complement(<1..3)
                     /locus_tag="GWK63_04415"
                     /pseudo
     CDS             complement(<1..3)
                     /locus_tag="GWK63_04415"
                     /inference="COORDINATES: similar to AA
                     sequence:RefSeq:WP_010336579.1"
                     /note="incomplete; partial on complete genome; missing
                     C-terminus; Derived by automated computational analysis
                     using gene prediction method: Protein Homology."
                     /pseudo
                     /codon_start=1
                     /transl_table=11
                     /product="IS110 family transposase"
```

Figure 5. Additional evidence supporting the JC affecting bcs operon IV in the parental strain. The reads covering the junction, when analysed by BLAST, show a clear association with transposase activity.

Table 1. Translation of $ccpA_x$ generated using Geneious. The 13 consecutive proline residues, where replication slippage events were detected, are highlighted.

| Original annotation | BLAST annotation | Genomic position |
|---|---|---|
| **dosP_1/Hypothetical protein** | Putative bifucntional DGC_PDE / EAL-domain containing protein | 438431-442444 |
| **dosP_2** | EAL domain containing protein | 768732-771215 |
| **putative signaling protein / dosp_3** | Putative bifucntional DGC_PDE / EAL-domain containing protein | 935004-939007 |
| **dosP_4** | EAL domain containing protein | 2393877-2396450 |
| **hypothetical protein/hypothetical protein** | EAL domain containing protein/ putative bifunctional DGC_PDE | 2400866-2404982 |
| **dosP_5** | EAL domain containing protein | 2889745-2891967 |
| **dosP_6** | Putative bifunctional DGC_PDE | 2892011-2894218 |
| **hypothetical protein** | EAL domain containing protein | 3602045-3604261 |
| **dosP_7/putative signaling protein** | EAL domain containing protein/ putative bifunctional DGC_PDE | 3604360-3608735 |

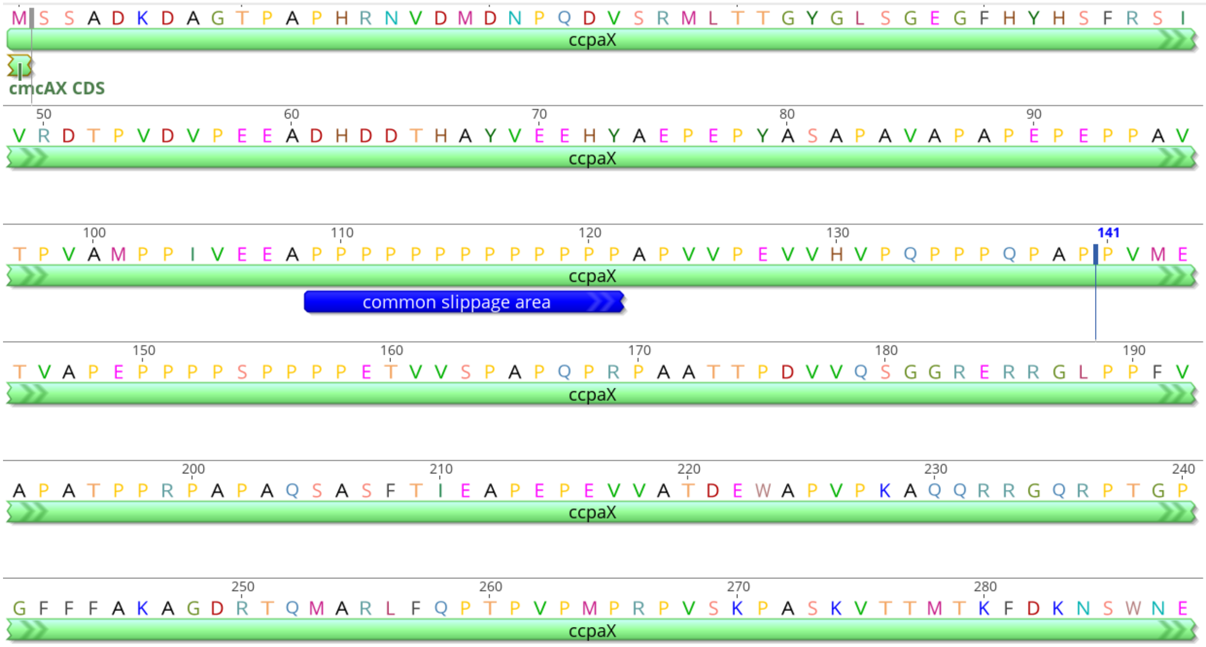


Figure 6. Translation of *ccpAx* made on Geneious. The image highlights the region containing 13 consecutive prolines, where replication slippages were detected in the parental strain; Flask A (rounds 4, 7, and 10); Flask B (rounds 2, 3, 5, 6, and 7); and Flask C (rounds 1, 2, and 10).

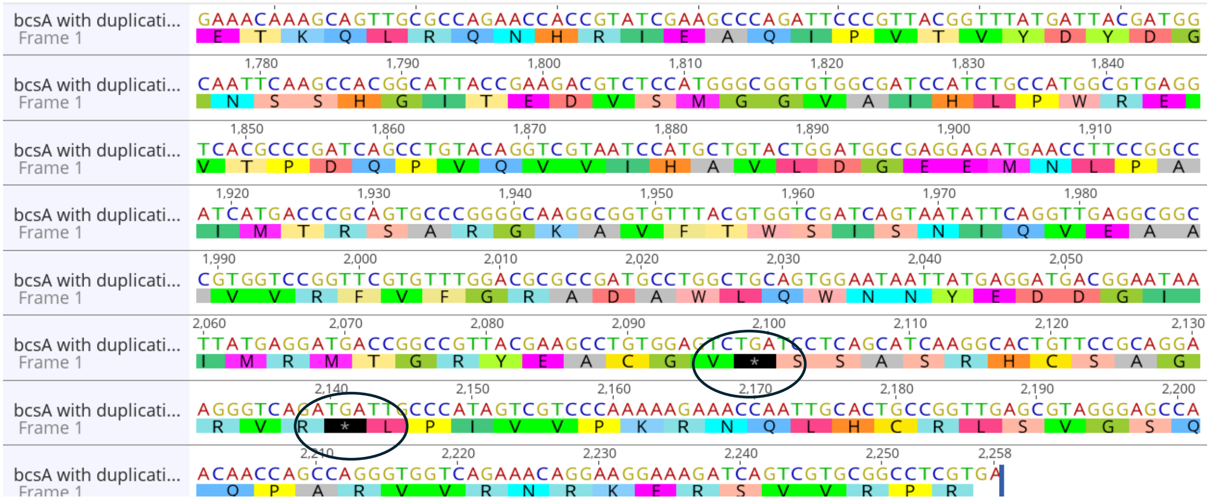


Figure 7. Translation, made on Geneious, of the *bcsA* with the 20 bp duplication that affected it in Flask C (rounds 9 and 10). The duplication leads to two premature stop codons.

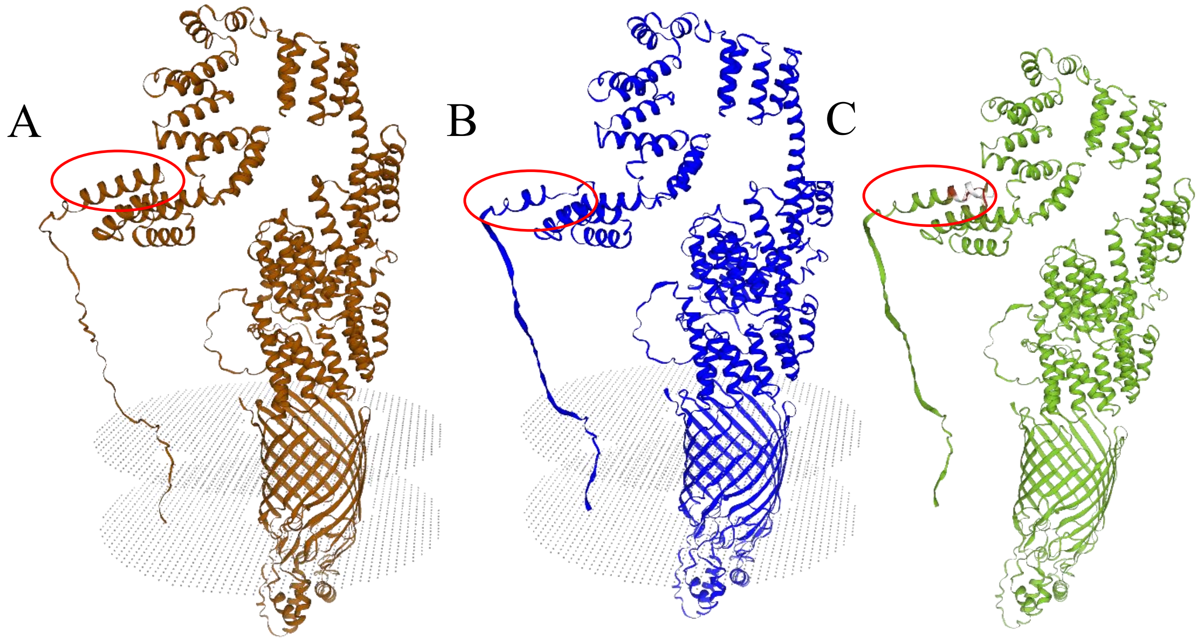


Figure 8. 3D structures ( made on SwissModel) of wild type *bcsC,* and the *bcsC* mutated by the 21 bp deletion that removes the ARFWLQQ amino acid sequence. A) The wild type *bcsC*. B) The mutated *bcsC*. C) superimposition of wild type bcsC 3D model on the on mutant bcsC 3D model. Green colour indicate a match between the two models, while the red color indicate a mismatch. For A,B,C the region affected by the mutation is circled in red.

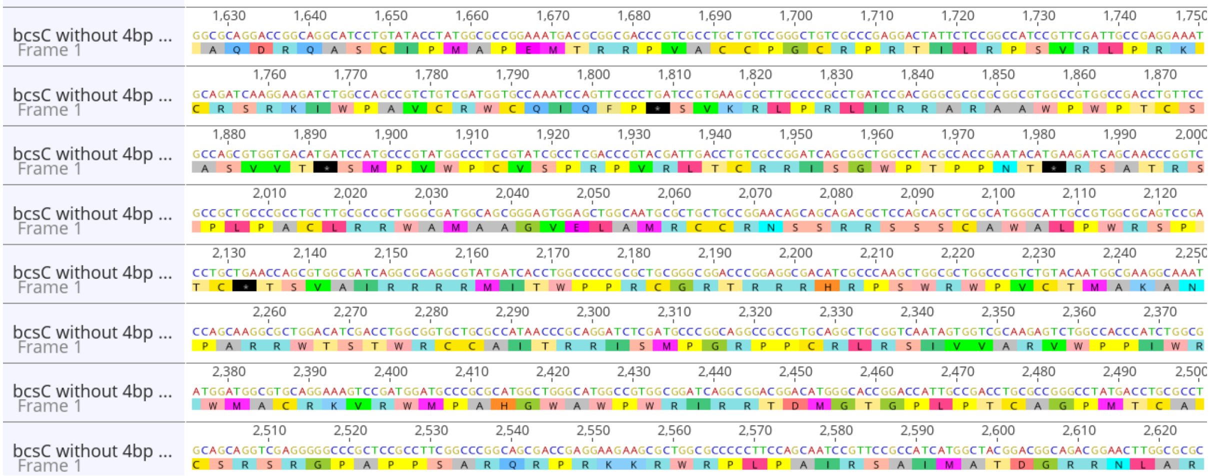


Figure 9. Translation, made on Geneious, of the *bcsC* region following the 4 bp deletion detected in Flask A ( round5), Flask B ( rounds3,4,5) and Flask C (rounds5,6,7,8). The deletion cause multiple premature stop codons (shown in black).

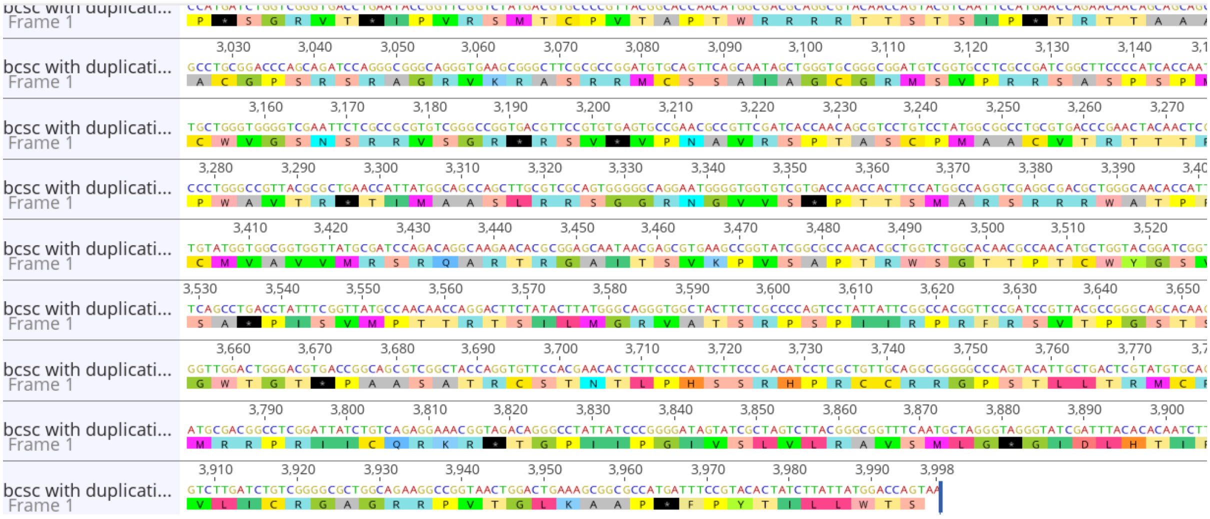


Figure 10. Translation, made on Geneious, of the *bcsC* after the the duplication event (bp 1517909-928), which cause several premature stop codons ( in black) in Flask C, round 6.

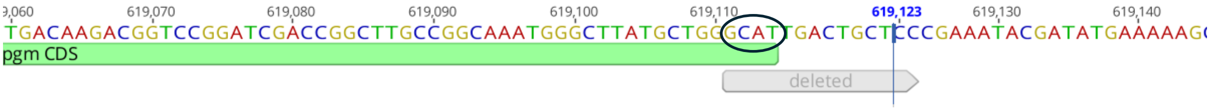


Figure 11. Visualization, made on Geneious, of the deletion causing the loss of the start codon in the *pgm* gene, which is located on the reverse strand. The mutation was detected in Flask C, rounds 8,9,10.

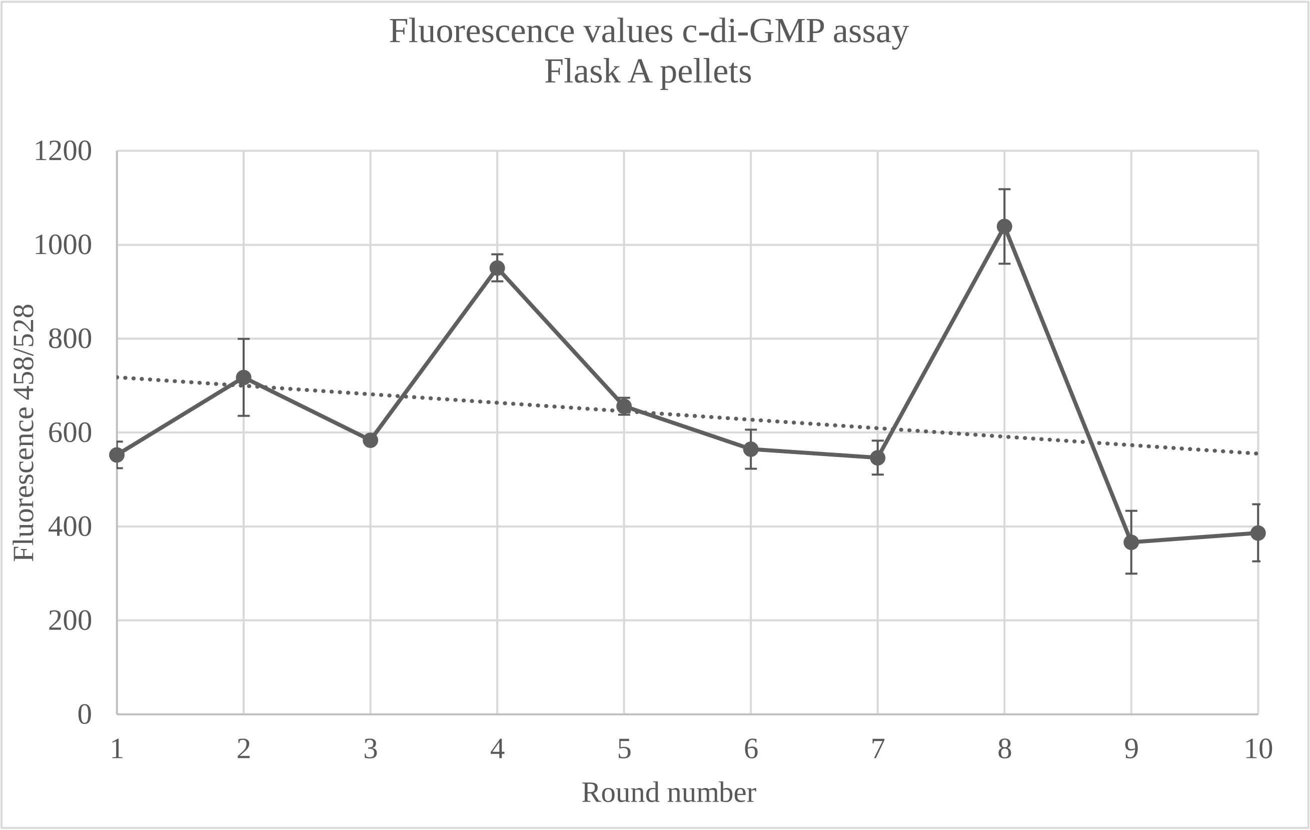


Figure 12. Fluorescence values from the intracellular c-di-GMP assay for pellets collected from Flask A after each cultivation round.
